# BFVD v3–UniProt-complete, improved viral protein structure predictions

**DOI:** 10.64898/2026.09.16.752260

**Authors:** Rachel Seongeun Kim, Olga Pimenova, Eli Levy Karin, Milot Mirdita, Martin Steinegger

## Abstract

The Big Fantastic Virus Database (BFVD) v3 is the most comprehensive resource for ColabFold-AlphaFold2-predicted viral protein structures. It holds 5,776,417 structures of nearly all viral sequences in UniProt 2025_03, a 16.4-fold increase compared to the representative-only catalogs of BFVD v1 and v2. Structure prediction quality has also improved, with high-confidence predictions accounting for 75.3% of BFVD v3 entries. This is due to Logan’s enormous sequence assembly, now mined for the entire BFVD v3 instead of only for shallow alignments, alongside continued prediction quality improvements. BFVD v3 covers 72.7% of ICTV’s virus species, 1.54-fold more than BFVD v2. With this version, the fraction of fully covered viral reference proteomes increases from 1.5% to 72.6%. BFVD v3 thus brings us closer to a structural catalog of the known virosphere and is freely available at bfvd.steineggerlab.workers.dev and bfvd.foldseek.com.

## INTRODUCTION

Following advances in protein structure prediction [1, 2], recent years have seen the emergence of dedicated viral structure prediction resources: the Big Fantastic Virus Database (BFVD) [3], Viral AlphaFold Database [4], Nomburg et al. [5] for eukaryotic viruses, Viro3D [6] for human and animal viruses, and Phold [7] for bacteriophages. This reflects the broader recognition that protein structures offer a route to studying viral function and evolution that sequence-based annotation alone cannot, given how rapidly viral sequences diverge [8, 9, 10, 11]. Among these, BFVD v1 comprised 351,242 predicted structures for the UniRef30 [12] cluster representatives of viral origin in UniProt 2023_02 [13], thereby complementing AlphaFold Protein Structure Database (AFDB) [14], which had excluded viral sequences. Since then we released BFVD v2, which improved the predictions of a subset of v1 entries using optimized AF2 parameters (Supplementary Results). BFVD v2 has been incorporated into AFDB, extending its structural coverage beyond the tree of life.

Structure prediction in BFVD v2 relied on AlphaFold2 (AF2) modeling with ColabFold [15], using MMseqs2 [16] to search against its default environmental database and construct multiple sequence alignments (base-MSAs). For sequences with shallow base-MSAs (< 30 homologs), prediction quality was boosted by mining additional homologs from the Logan assembly [17] of the Sequence Read Archive [18]. The v2 prediction setup included the early stop condition, which saves improvement steps once the predicted Local Distance Difference Test (pLDDT) score reaches 85.

The use of UniRef30’s viral cluster representatives in BFVD v2 has two main limitations. First, representing a cluster with a single structure discards the structural diversity among its members, obscuring how a protein’s fold diverges across closely related sequences. Second, a UniRef30 cluster representative was included in BFVD v2 only if it was a viral protein, thereby missing viral sequences whose representative was a non-viral sequence.

Here we present BFVD v3, which provides 5.8M predicted structures for nearly all viral sequences in UniProt 2025_03 [19], with Logan homology mining applied across the entire set instead of only to sequences with a shallow base-MSA and prediction run for three recycles without early stopping. BFVD v3 faithfully reflects UniProt’s taxonomic composition rather than the skewed representation by cluster representatives, and it covers 72.7% of the International Committee on Taxonomy of Viruses (ICTV)’s [20] virus species, a 1.54-fold improvement over BFVD v2.

## MATERIALS AND METHODS

### Acquisition of sequences and MSA generation

UniProt 2025_03 [19], which was used for the AFDB v6 release [14], was downloaded and 5,841,128 sequences assigned a taxonomic identifier (“taxid”) descending from taxid 10239 (“Viruses”) were retained. Sequences longer than 2,000 residues were excluded, retaining 5,776,417 (98.9%) of viral proteins in UniProt 2025_03.

For each sequence, a multiple sequence alignment (base-MSA) was obtained using *colabfold_search* (MMseqs2 commit 6f4523, sensitivity *−s* 7) against the reference databases `uniref30_2302_db` and `colabfold_envdb` `_202108_db`.

To augment each base-MSA with additional homologs, we used the released Logan50 resource, a 50% sequence identity clustering of the Logan assembly, excluding human-derived sequences and comprising 3,009,464,616 sequences. We queried Logan50 using MMseqs2 *search* (version 18.8cc5c), with parameters: *−e* 0.1, −−max-seqs 1000 and two search iterations, followed by *result2msa* to construct a Logan50-MSA, which was appended to the base-MSA. For each of the base- and Logan50-MSA, we computed the effective number of sequences (*N*_eff_) using hhmake [21] (version 3.3.0).

### Structure prediction

Before predicting the full v3 dataset, we benchmarked prediction accuracy under four prediction settings on a small subset of experimentally resolved protein structures (“ground truth”) that were not seen by AlphaFold2 (AF2) during training (full details in Supplementary Results). We then used the best-performing settings to predict all v3 structures from the Logan50-appended MSAs using ColabFold-AF2 (version 1.6.0), with a single fixed AF2 model (model 4), three recycles, and no template. For low-confidence predictions (ColabFold-AF2 pLDDT *<* 70) whose length was *<* 450, we ran ESMFold+ProteinTTT_MSA_ [22] and used its predictions for 40,524 proteins (0.7% of v3 entries) that met our confidence and improvement criteria (see Supplementary Results). For the analysis of v3 pLDDT we used the values computed by ESMFold+ProteinTTT_MSA_ for the replaced entries and otherwise, the ColabFold-AF2 values.

### Proteome and taxonomic coverage

We downloaded UniProt’s viral reference proteomes (release 2026_02) and used the proteins labeled as “canonical sequences” for each reference proteome. We searched these sequences against BFVD v2 and v3 separately (MMseqs2, version 18.8cc5c, −−seq-id 1.0, *−*c 1.0), counting a protein as covered by v2 (v3) if it had an exact sequence match in v2 (v3). Per-proteome coverage is the fraction of a proteome’s proteins with such a match.

Each BFVD entry (v2 and v3) is associated with an NCBI taxid through its UniProt identifier. We mapped BFVD taxids to ICTV-classified species [20] as follows. Each ICTV species is associated with GenBank accession(s), which are linked to NCBI taxid(s) at the species level. We then looked up each BFVD taxid in the list of taxids associated with ICTV species. If we could not find the BFVD taxid in the list, we looked for an exact match between the taxon name and an ICTV species name. The rest remained unresolved. At each taxonomic rank, coverage is the fraction of ICTV-recognized taxa with at least one mapped BFVD entry.

### Sequence and structural clustering

To compare the pLDDT distributions of BFVD v3 and v2, we clustered v3 by its sequences (MMseqs2 clustering, minimum sequence identity 0.3, coverage 0.9), to resemble v2, which was constructed from sequence cluster representatives. To examine sequence and structure variance, we jointly clustered v2 and v3 twice. Once by the entries’ sequences (MMseqs2, version 18.8cc5c, 30% sequence identity, 90% coverage) and once by their structures (Foldseek, version 10.941cd33, *easy-cluster*, 70% coverage). In each clustering, we classified clusters as “v2-unique”, “v3-unique”, or “shared”, depending on the origin of their members; we also labeled clusters as “singleton” (single cluster member) or “non-singleton”.

### Webserver

To enable interactive exploration of BFVD v3, we extended the BFVD v2 webserver [3], available at bfvd.foldseek.com.

Each of v3’s 5.8M entries has a dedicated page annotated with its UniProt proteome (44.3% of entries), the ICTV identifiers mapped in this study (Materials and Methods), and a host organism from UniProt or, where absent, ICTV’s coarse host category (75.4% of entries). For each entry we also display its v3 sequence cluster (Materials and Methods) with the cluster’s members, summary statistics, and taxonomic distribution as a Sankey diagram (code adapted from [23]). For navigation to structurally similar entries, we searched BFVD v3 against itself with Foldseek [24] (version 10.941cd33, *−s* 9.5, *−e* 0.01); 98.0% of entries have at least one hit, listed on their page.

We display structures with Mol* [25], rebuilding full-atom models from the stored C*α* traces with PULCHRA [26], and superposing an entry onto its sequence cluster members or structural neighbors with TM-align [27]. BFVD can be queried by UniProt accession, by taxonomic name or identifier, or by structure, using Foldseek search.

## RESULTS

### Scale and taxonomic coverage of BFVD v3

BFVD v3 comprises 5,776,417 predicted structures – a 16.4-fold increase over BFVD v2’s 351,242 structures, and a near-complete (98.9%) coverage of the viral sequences in UniProt 2025_03 (**Figure 1a**).

**Figure 1.**
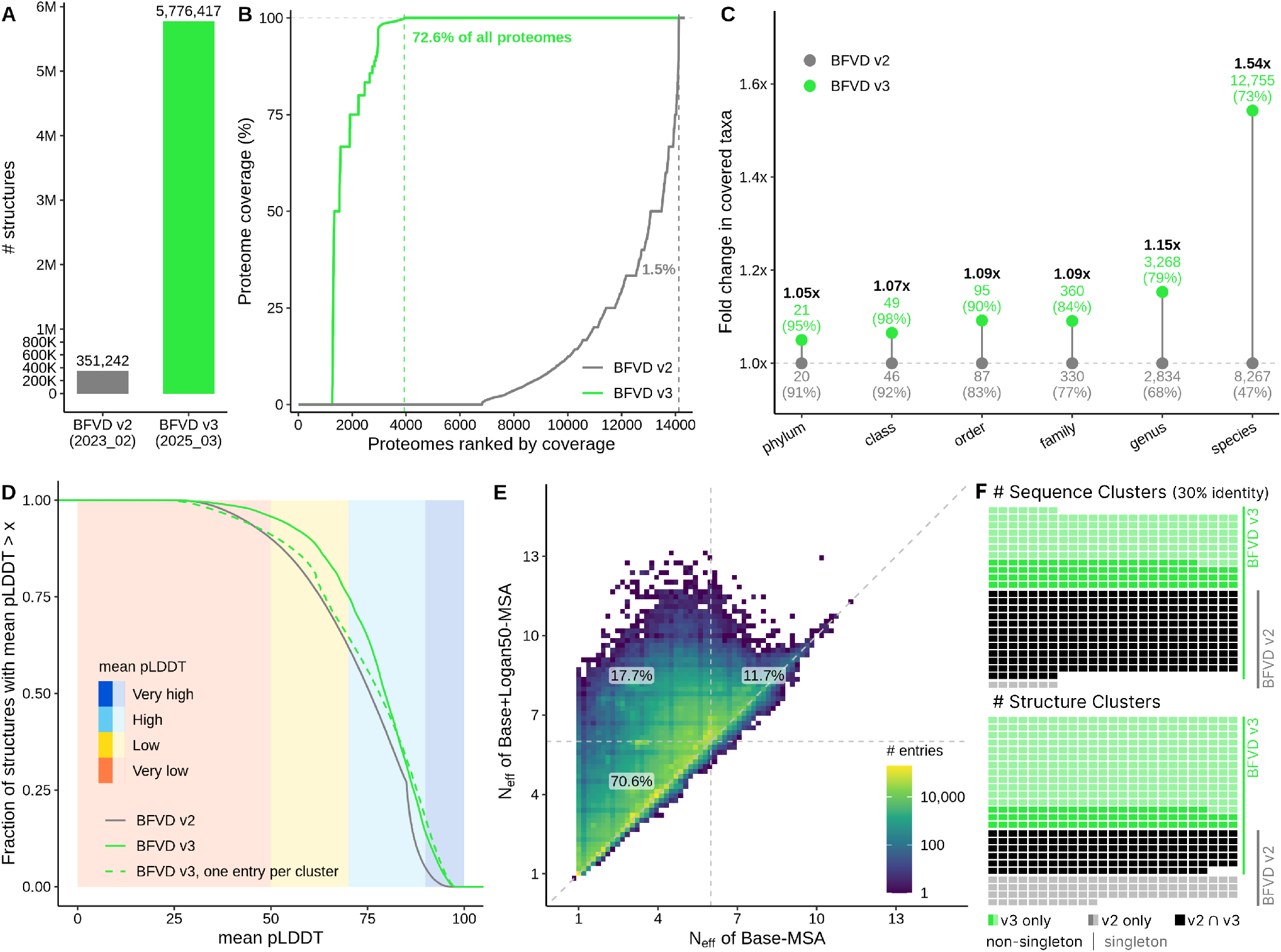
BFVD v3 vs. v2 in terms of scale, taxonomic coverage, prediction quality and alignment depth. **(a)** Number of deposited structure predictions. **(b)** Over 14,000 viral reference proteomes (x-axis) ranked by their coverage (y-axis), computed for each from the fraction of its proteins with an exact sequence match in v2 (gray) and v3 (green). **(c)** Fold change coverage of ICTV taxa at ranks phylum through species. At each rank, points are labeled with the number of taxa, for which at least one structure is included in v2 (gray) and v3 (green); percent coverage out of total ICTV labels for the rank is given in parentheses. Along the y-axis, the coverage by v2 is a baseline at 1.0 with v3 points placed at the fold change values (black labels). **(d)** Cumulative mean pLDDT distributions. Two curves depict v3: the scores of all entries (green, solid) and the scores of representatives following clustering at 30% sequence identity, to resemble v2 construction conditions (green, dashed); v2 is shown in gray. Background shading marks standard confidence ranges. **(e)** Effective sequence number (*N*_eff_) before (x-axis) and after Logan50 mining (y-axis) as a 2D density plot. Color indicates the number of entries at each point. Dashed lines at *N*_eff_=6 mark the boundary for high effective diversity; Logan50 mining moves 17.7% of v3 entries from below this threshold to at or above it. **(f)** Composition of joint (v2 + v3) sequence (top) and structure (bottom) clusters (1,200 clusters per cell). Colors indicate v2-uniqueness (gray), v3-uniqueness (green), or shared (black); lighter gray and green denote singleton clusters. BFVD v2-unique non-singleton clusters are not visible at this scale (n = 175 for sequence, n = 521 for structure).

Considering a sequence in a proteome as covered if it exactly matches a BFVD sequence, we find that v3 fully covers 72.6% of viral reference proteomes compared to 1.5% by v2 (**Figure 1b**). Furthermore, the fraction of proteomes with no structural coverage at all falls from 47.6% to 8.8% between the versions.

We next mapped every BFVD v2 and v3 entry to the ICTV’s Virus Metadata Resource [20] to quantify taxonomic coverage. While ICTV’s higher ranks were already fully covered in BFVD v2 (10/10 realms, 11/11 kingdoms), coverage improves from v2 to v3 at every lower rank. At the species level, v3 covers 72.7% of ICTV’s 17,554 cataloged species, compared to 47.1% in v2, a 1.54-fold increase (**Figure 1c**). Consistent with this finding but independently of ICTV, counting distinct NCBI taxids in v2 and v3 reveals a 2.12-fold increase in unique species-level taxa (20,847 to 44,124) from v2 to v3.

The reliance of BFVD v2 on UniRef30 representatives introduced a taxonomic bias that did not reflect the underlying distribution of viral sequences in UniProt; for example, 62.3% of the structures in v2 belong to the kingdom Heunggongvirae, compared to only 29.33% in UniProt 2023_02. By predicting structures for individual UniProt sequences, BFVD v3 faithfully mirrors UniProt’s taxonomic composition, expanding coverage of previously underrepresented groups, such as RNA viruses.

### Prediction confidence and Logan50 homologs

Over 75% of BFVD v3 structures (4,351,255 predictions) are of high confidence (pLDDT ≥ 70), compared to 62.1% in v2 (**Figure 1d**, solid lines). However, the raw pLDDT distribution of BFVD v3 could be skewed simply by including multiple closely related sequences that are each predicted with high confidence. To eliminate this effect when comparing to v2, we clustered the sequences of v3 to resemble the use of representatives by v2 (Methods). The clustered-v3 pLDDT curve closely tracks that of v2 until a pLDDT value of about 75, but improves over v2 for values greater than that (**Figure 1d**, dashed green line). This means v3 adds a small fraction of very high confidence structures that are not due to sequence redundancy. One factor that contributes to this high confidence fraction is the removal of early stopping, which curtailed the number of highly scoring structures in v2, as reported in [4].

The second factor is the extension of Logan homology mining to the entire dataset. In BFVD v2, Logan mining was applied to entries with a shallow base-MSA (about half the database), improving the pLDDT of 71.1% of them [3]. In BFVD v3, we extended this approach using Logan50, a clustering of Logan sequences at 50% sequence identity [17], applied to every entry regardless of base-MSA depth. Before Logan50 mining, 14.1% of BFVD v3 entries (811,890) had a shallow base-MSA. For these, Logan50 increased the median number of homologs from 8 to 15, moving 24.8% (200,967) above the shallow threshold.

Since structure prediction with AF2 is more directly affected by the effective sequence number (*N*_eff_), a measure of the MSA diversity [1], we measured it before and after Logan50 mining (**Figure 1e**). Across all v3 entries, median *N*_eff_ rose from 4.1 to 4.7 after adding Logan50 homologs (mean 4.0 to 4.6). For the shallow 14.1%, median *N*_eff_ rose from 1.3 to 1.8. At the boundary of high effective diversity (*N*_eff_=6), Logan50 mining raised 17.7% of entries from below this threshold to at or above it. This shows that Logan50 mining delivers genuine gains in evolutionary information.

### Growth in sequence and structural diversity

To explore the sequence and structure variance of BFVD v3 with respect to v2, we jointly clustered the databases (Methods). At 30% sequence identity clustering, 47.0% of all clusters (308,281) are v3-unique, 51.6% (338,622) are shared with v2, and only 1.4% (8,924) are v2-unique. This shows that v3 both nearly covers v2’s sequence space and doubles it to include sequences added to UniProt since v2 and sequences whose UniRef30 cluster representative is not viral.

Structurally, 61.5% (450,113) of all clusters are v3-unique, 24.2% (176,990) are shared and 14.2% (104,204) are v2-unique. Since singleton clusters represent weaker evidence for structural validity, we examined their prevalence among v2- and v3-unique clusters. Nearly all v2-unique structural clusters (99.5%) and most v3-unique clusters (80.8%) were singletons (**Figure 1f**, bottom), consistent with the known difficulties of clustering and structural modeling of viral proteins [3]. A closer inspection of v2-unique clusters further suggests these are not reliably modeled structures overlooked by v3: 76.1% are shorter than 100 residues and 55.8% have fewer than 30 sequence homologs.

Taken together, BFVD v3 captures 1.9-fold more sequence variance and 2.2-fold more structural variance than BFVD v2, better capturing the variance of the virosphere.

## DISCUSSION

BFVD v3 extends the resource introduced with BFVD v2 from a subset of viral sequence space to near-complete coverage of viral UniProt. This narrows the long-standing gap between sequence and structure coverage for viruses. By extending Logan mining to every entry in v3, we push viral homology detection much closer to what sequence databases can offer.

Unlike narrower viral structure resources, BFVD v3 holds a broad yet detailed image of the known virosphere, thereby enabling cross-lineage comparative analyses. However, since UniProt carries a sampling bias that does not uniformly reflect the virosphere, BFVD v3 inherits it: Influenza A virus alone, for instance, accounts for 13.3% of UniProt 2025_03’s viral sequences. This bias and the remaining ICTV species-level gap in BFVD’s coverage (missing 27%) could be reduced in a future update of BFVD, when new viral genomes become available. Finally, to limit the computational demand of structure prediction we excluded sequences over 2,000 residues (1.1% of viral UniProt). This fraction could have increased the ICTV species coverage by an additional 1.3%. With its advancements, BFVD v3 positions the community to move from cataloging viral sequences to understanding their structural biology at scale.

## Supporting information

Supplementary Results

## CODE & DATA AVAILABILITY

All metadata and predicted structures, accompanying Foldseek database, can be freely downloaded from bfvd.steineggerlab.workers.dev. BFVD v3 can be browsed at bfvd.foldseek.com. The webserver code is available at github.com/rachelse/bfvd3-web.

## ACKNOWLEDGEMENTS

We thank Anton Bushuiev and Josef Sivic for their advice on integrating ESMFold+ProteinTTT_MSA_, and NVIDIA for providing computational resources for structure prediction.

