## Supplementary Results for "BFVD v3–UniProt-complete, improved viral protein structure predictions"

### Supplementary Materials for BFVD v3–UniProt-complete, improved viral protein structure predictions

#### SUPPLEMENTARY RESULTS

##### From BFVD v1 to v2: prediction refinements

BFVD v1 predicted structures for 351,242 UniRef30 viral cluster representatives using ColabFold-AF2 with 3 models and 3 recycles each, retaining the highest-ranked model per entry. For the 175,788 entries with shallow base-MSAs, homologs from the Logan assembly were mined to augment the alignments.

In BFVD v2, we revisited these shallow-MSA entries with an additional 12-recycle prediction on the Logan-augmented MSA, generating up to three prediction settings per entry: base-MSA with 3 recycles, Logan-augmented MSA with 3 recycles, and Logan-augmented MSA with 12 recycles. For each setting, 3 models were run, and the overall highest-confidence prediction (by pLDDT) across all settings and models was retained as the v2 entry. The remaining 175,454 entries with sufficient base-MSA homologs were carried over unchanged from v1.

##### Benchmark to decide structure prediction conditions

We evaluated two strategies to improve structure prediction: augmenting MSAs with Logan50 homologs, and using BFVD v2's predicted structures as templates. For benchmarking, we collected ground-truth structures of viral proteins not seen by AlphaFold2 (AF2) [1] during training as follows.

1. Starting from 4,757 viral UniProt entries mappable to Protein Data Bank (PDB) [2] structures, we kept those whose earliest deposition date is after AF2's training cutoff (30 April 2018), thereby retaining 2,140.
2. Of these we kept entries of length $\leq$1,500 residues whose mapped PDB chain covers $>$0.9 of the UniProt sequence, retaining 1,172.
3. We then removed any entry whose sequence matched that of a pre-cutoff PDB deposition (MMseqs2 commit 6f4523, sequence identity $>$0.7, coverage $>$0.9), retaining 955 entries.
4. Finally, we reduced their redundancy by MMseqs2 clustering (minimum sequence identity 0.7, coverage 0.8), yielding 821 representative sequences, each paired with one ground-truth PDB chain of highest coverage.

For each of the 821 sequences in our benchmark we constructed a base-MSA and a Logan50-MSA as described in Materials and Methods. Additionally, for each sequence we mined templates using *colabfold_search* (MMseqs2 commit 6f4523, sensitivity $-s$ 7) against BFVD v2 (`bfvd_2023_02_v2`).

Each of the 821 sequences was predicted under all four combinations of prediction conditions, using only AF2 models 1 and 2 for template-based predictions. We assessed prediction confidence and structural alignment quality to the ground-truth structure, by LDDT [3] and TM-score [4], computed with Foldseek [5] (version 10.941cd33) *easy-search* (exact TM-score, exhaustive search). Under all four settings, the predictions of 53 sequences could not be aligned to their corresponding ground-truth structures and we set their LDDT/TM-score to $-0.1$.

Compared to the baseline setting (base-MSA, no template), Logan50-augmented MSAs improved LDDT for 60.5% of the proteins (TM-score for 54.9%), raising the mean LDDT from 0.701 to 0.718 (mean TM-score from 0.566 to 0.582). Of all four settings, Logan50-augmented MSAs without a template achieved the highest accuracy in both metrics (LDDT shown in **Supp. Figure 1a, b**). Adding Logan50 to a setting also consistently translated into gains in prediction confidence (pLDDT, **Supp. Figure 1c**, left pair and right pair).

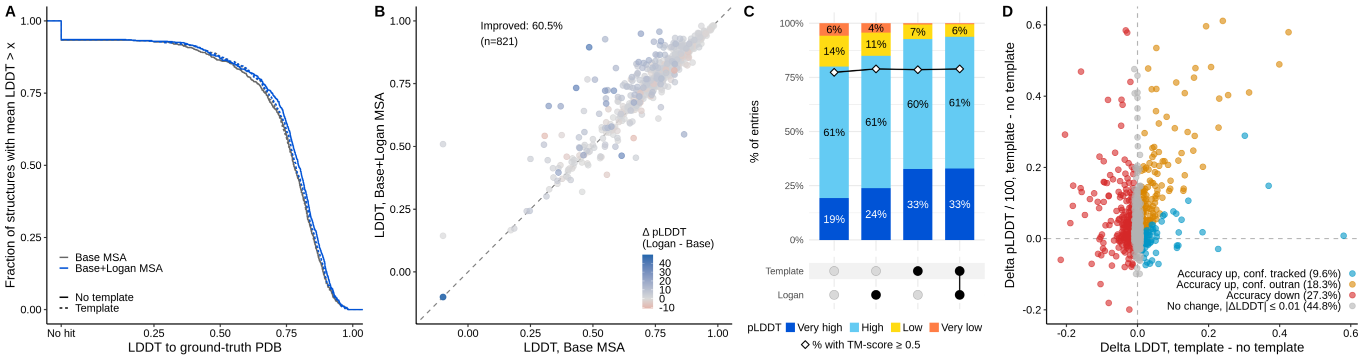


**Supplemental Figure 1. Prediction conditions benchmark.** Each prediction under four tested conditions was aligned to its corresponding ground-truth PDB structure. Alignment LDDT and prediction confidence (pLDDT) were recorded for 821 benchmark proteins under each condition. **(a)** LDDT to ground truth (x-axis) and the fraction of structures scoring above it (y-axis). Each tested condition is a curve. **(b)** Per-entry LDDT for base-MSA (x-axis) and after Logan50-augmentation (y-axis). **(c)** Fraction of proteins in each pLDDT range for the four conditions. The fraction of accurately predicted proteins (TM-score $\geq 0.5$) is overlaid as a black diamond line. **(d)** Confidence change as a function of accuracy change. The LDDT (accuracy) of each prediction from the base-MSA (no template) condition was subtracted from the LDDT when predicting with a template (delta LDDT, x-axis) and similarly for the confidence measure pLDDT (delta pLDDT, y-axis). Colors: gray, $|\text{delta LDDT}| \leq 0.01$, negligible accuracy change; red, delta LDDT $< -0.01$ (template reduces accuracy); yellow, pLDDT $>$delta LDDT $>0.01$ (template increases accuracy and is overconfident); and blue, pLDDT $<=$delta LDDT $>0.01$ (template increases accuracy, confidance matches).

Structural templates, by contrast, often raised prediction confidence without improving their accuracy. On top of base-MSAs, templates increased the fraction of predictions with mean pLDDT $\geq 70$ from 80.0% to 92.7% and almost eliminated the “Very low” pLDDT category (**Supp. Figure 1c**, 1st and 3rd bars), while LDDT increased for only 45.1% of the proteins with a marginal gain (mean +0.04) and a similar-sized loss among the rest (mean -0.03). Examining each protein by the template effect, only 9.6% gained accuracy without overconfidence, while 18.3% were overconfident and 27.3% lost accuracy (**Supp. Figure 1d**). Adding templates on top of Logan50-augmentation showed a similar trend: the mean LDDT decreased from 0.718 to 0.711 with templates (**Supp. Figure 1a**, blue solid and dashed lines), while pLDDT values increased (**Supp. Figure 1c**, 2nd and 4th bars). We therefore opted not to use templates for the full v3 set.

#### Rescue of low-confidence predictions with ESMFold+ProteinTTT$_{\text{MSA}}$

While ColabFold-AF2 predicted most BFVD v3 structures with high confidence, 25% had pLDDT $< 70$. In the following, we describe our procedure to improve the prediction of a subset of them with ESMFold [6] enhanced by ProteinTTT$_{\text{MSA}}$ [7], which together complement ColabFold-AF2. Specifically, we focused on 790,996 low-pLDDT proteins that were shorter than 450 residues. This length cutoff was set by the GPU memory available for running ESMFold+ProteinTTT$_{\text{MSA}}$ on our cluster.

ProteinTTT fine-tunes a pre-trained protein language model, like ESM2, to a specific protein sequence by minimizing the masked language modeling loss on its sequence, improving downstream predictions. ESMFold+ProteinTTT$_{\text{MSA}}$ adapts the ESM2 backbone to all homologs in an MSA, rather than to a single sequence. This allows the model to leverage evolutionary information from related sequences, without labeled data or retraining.

*Prediction setting with ESMFold+ProteinTTT*$_{\text{MSA}}$. We ran ESMFold+ProteinTTT$_{\text{MSA}}$ on the 790,996 candidate proteins using the Logan50-appended MSAs that were used for ColabFold-AF2 (Materials and Methods), with the following hyperparameters: 20 optimization steps, learning rate 0.04, MSA sampling strategy `neighbors`, gradient clipping max norm 1.5, LoRA rank 64, LoRA $\alpha$ 128, 64 gradient accumulation steps and mask ratio 0.2. To downweight redundant homologs, we sampled sequences with probability inversely proportional to their neighborhood size (within 0.2 normalized Hamming distance; 80% identity), following [8].

*Calibrating the pLDDT of ESMFold+ProteinTTT*$_{\text{MSA}}$. To enable direct comparison with ColabFold-AF2's pLDDT, we recalibrated ESMFold+ProteinTTT$_{\text{MSA}}$ confidence score using logit Platt scaling:

$$\widehat{\text{pLDDT}} = 100\sigma\left(a\,\text{logit}\left(\frac{\text{pLDDT}}{100}\right) + b\right), \qquad \textbf{(1)}$$

where $\sigma$ is the logistic function and $\text{logit}(x) = \ln\frac{x}{1-x}$. We fit the parameters $a = 0.5376$ and $b = -0.1063$ by cross-entropy on 16,051 per-residue $(\text{pLDDT}, \text{LDDT})$ pairs from 106 BFVD v3 benchmark proteins with a PDB structure (out of 821, see section above), selected by requiring ColabFold-AF2 pLDDT $< 70$ and length $< 450$. This calibration preserves the $[0, 100]$ range and reduces the mean per-residue pLDDT bias relative to true LDDT from $+6.6$ to approximately 0.0.

*Criteria for prediction replacement.* We set out to exclude ESMFold+ProteinTTT$_{\text{MSA}}$ predictions with extended helical structures, which are known to have inflated pLDDTs. To do so, we computed the long-range contacts of each prediction, defined as residue pairs whose C$\alpha$ atoms are within 8 Å but separated by more than 24 residues in sequence. Proteins with no long-range contacts (typically extended helical structures) were excluded, leaving 565,069 predictions (71.4%). For the final BFVD v3, we replaced 40,524 ColabFold-AF2 predictions (0.7% of v3) with ESMFold+ProteinTTT$_{\text{MSA}}$

ones whose calibrated pLDDT was greater than 60 and at least 10 points higher than ColabFold-AF2.

*Tuning hyperparameters.* We tuned on 100 BFVD v3 proteins with ColabFold-AF2 pLDDT $>90$, ESMFold pLDDT $<70$ and length $<450$ residues, i.e. cases in which ESMFold alone is unreliable but a confident ColabFold-AF2 prediction exists as a reference. A cyclic coordinate search scored by mean pLDDT over three seeds selected: learning rate $0.04$ with a constant schedule, 64 gradient accumulation steps, LoRA $(r,\alpha)=(64,128)$, 20 optimization steps, gradient clipping at max norm $1.5$, mask ratio $0.2$, and inverse-degree sub-MSA sampling at $80\%$ identity. Within the ranges searched, the MSA-sampling strategy affected pLDDT more than any single hyperparameter (84.4 for inverse-degree weighting vs. 83.3 uniform, 79.3 DBSCAN partitioning [9], 74.4 original alignment order). Gradient clipping is required: without it, ESMFold+ProteinTTT$_{\mathrm{MSA}}$ diverges on part of the tuning set until the structure module returns no residues and the prediction fails.
